## Supplementary Figures 1-27 for "Differentiating mechanism from outcome for ancestry-assortative mating in admixed human populations"

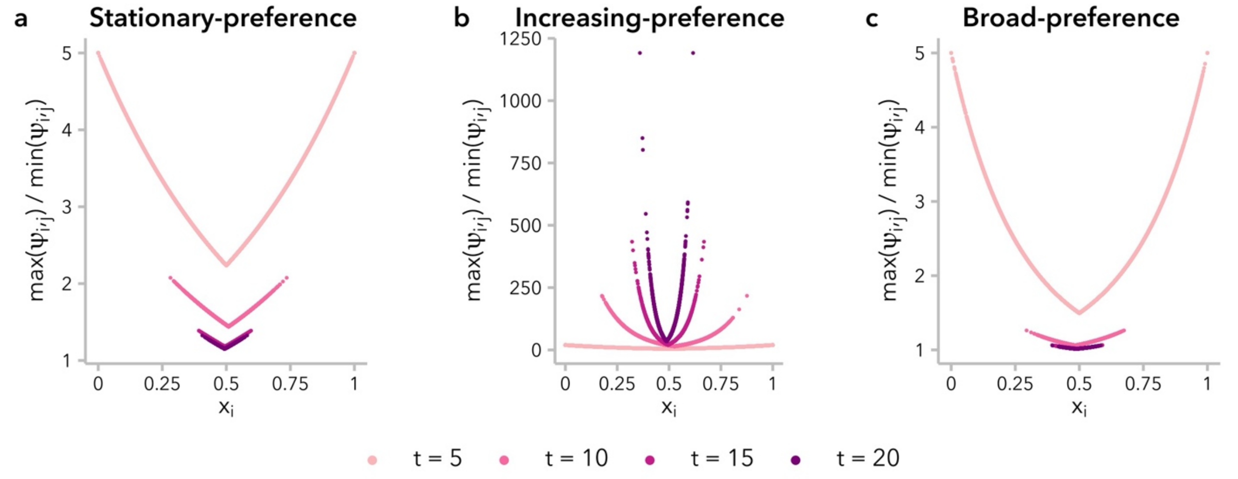


**Figure S1.** Individuals differ in "choosiness" – quantified as the ratio of $max(\psi_{i,j})\,/min\,(\psi_{i,j})$ – based on their own global ancestry proportion, $x_i$, and on the relative homogeneity of potential mates. $\alpha = 5$ is shown for the three models of non-random mating. **(a)** Under the stationary-preference model, $max(\psi_{i,j})\,/min\,(\psi_{i,j})$ approaches 1 (*i.e.,* random mating) for all values of $x_i$ over time. **(b)** Under the increasing-preference model, $max(\psi_{i,j})\,/min\,(\psi_{i,j})$ increases exponentially over each successive generation, resulting in increased choosiness over time. **(c)** Under the broad-preference model, $max(\psi_{i,j})\,/min\,(\psi_{i,j})$ approaches 1 for all values of $x_i$ more quickly than under the stationary-preference model.

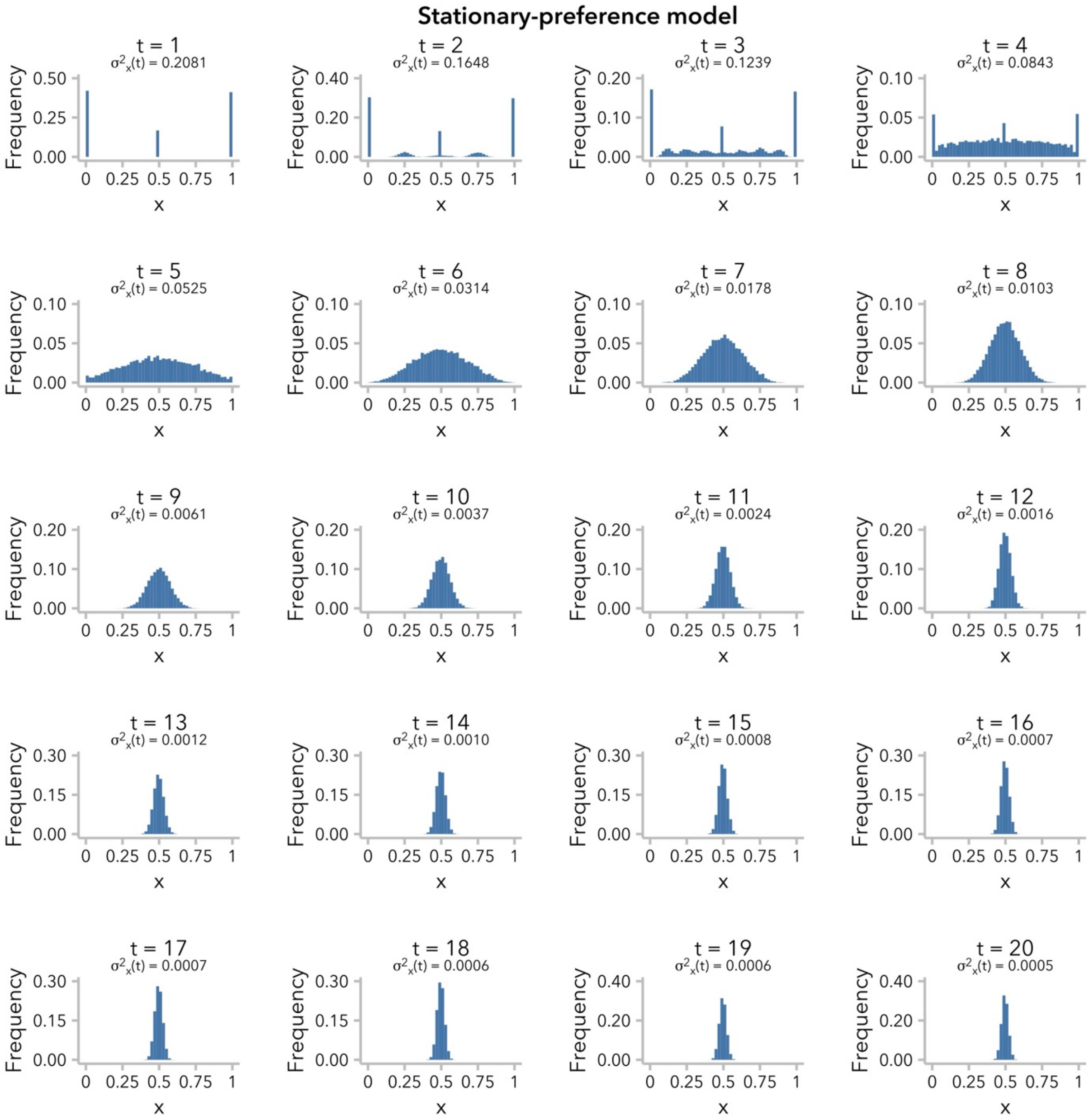


**Figure S2.** Under the stationary-preference model, variance in global ancestry proportion across individuals, $\sigma_x^2(t)$, decreases as admixture proceeds. The distribution of global ancestry proportion, $x$, is plotted for the first 20 generations post-admixture for $\alpha = 5$.

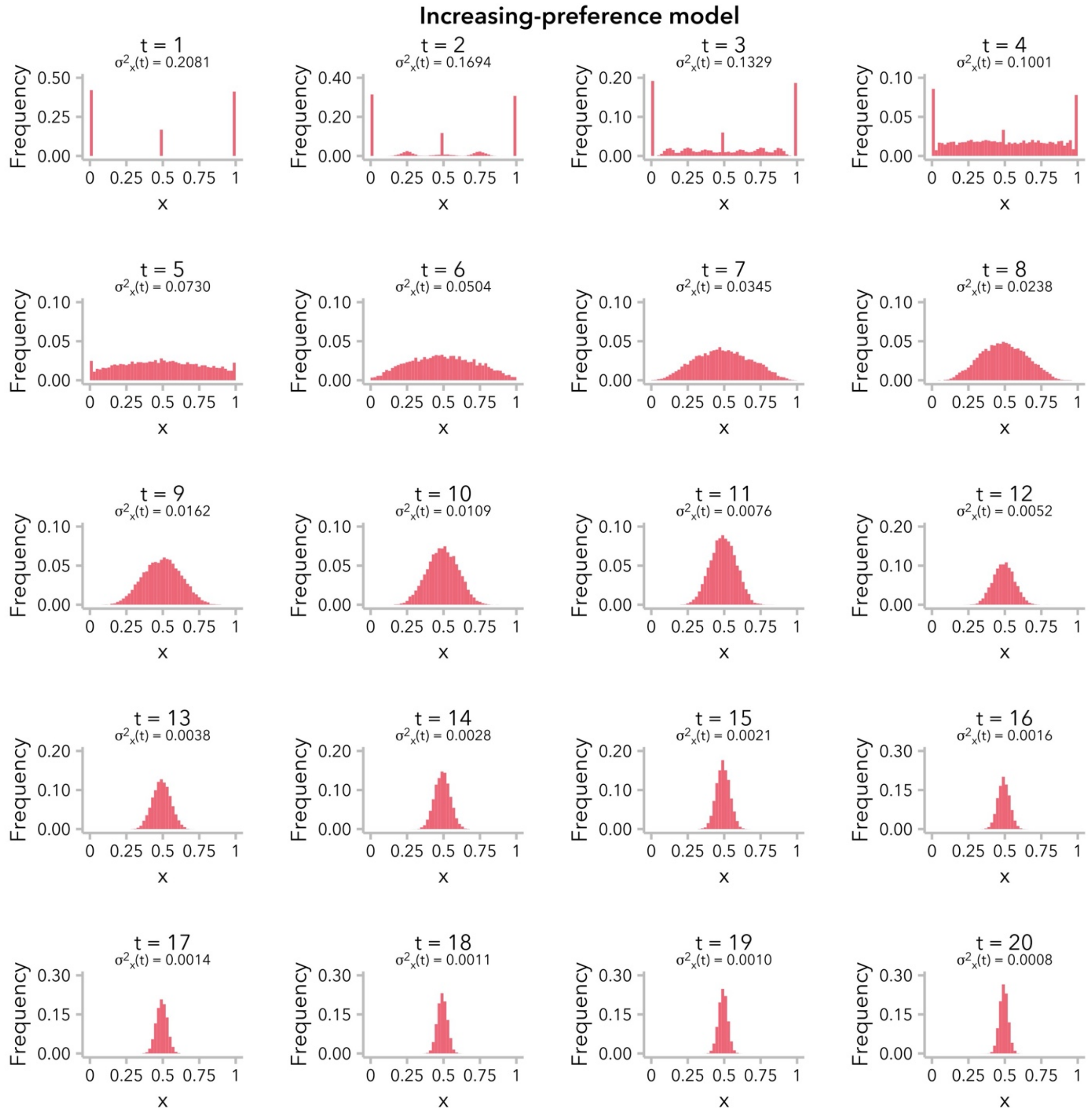


**Figure S3.** Under the increasing-preference model, variance in global ancestry proportion across individuals, $\sigma_x^2(t)$, decreases as admixture proceeds. The distribution of global ancestry proportion, $x$, is plotted for the first 20 generations post-admixture for $\alpha = 5$. To compensate for the decreased $\sigma_x^2(t)$ over time, the mate-choice parameter $c$ is scaled by $\sigma_x^2(t)$, meaning that the ratio of $max(\psi)\ /min\ (\psi)$ increases over time (see **Figure S1b**).

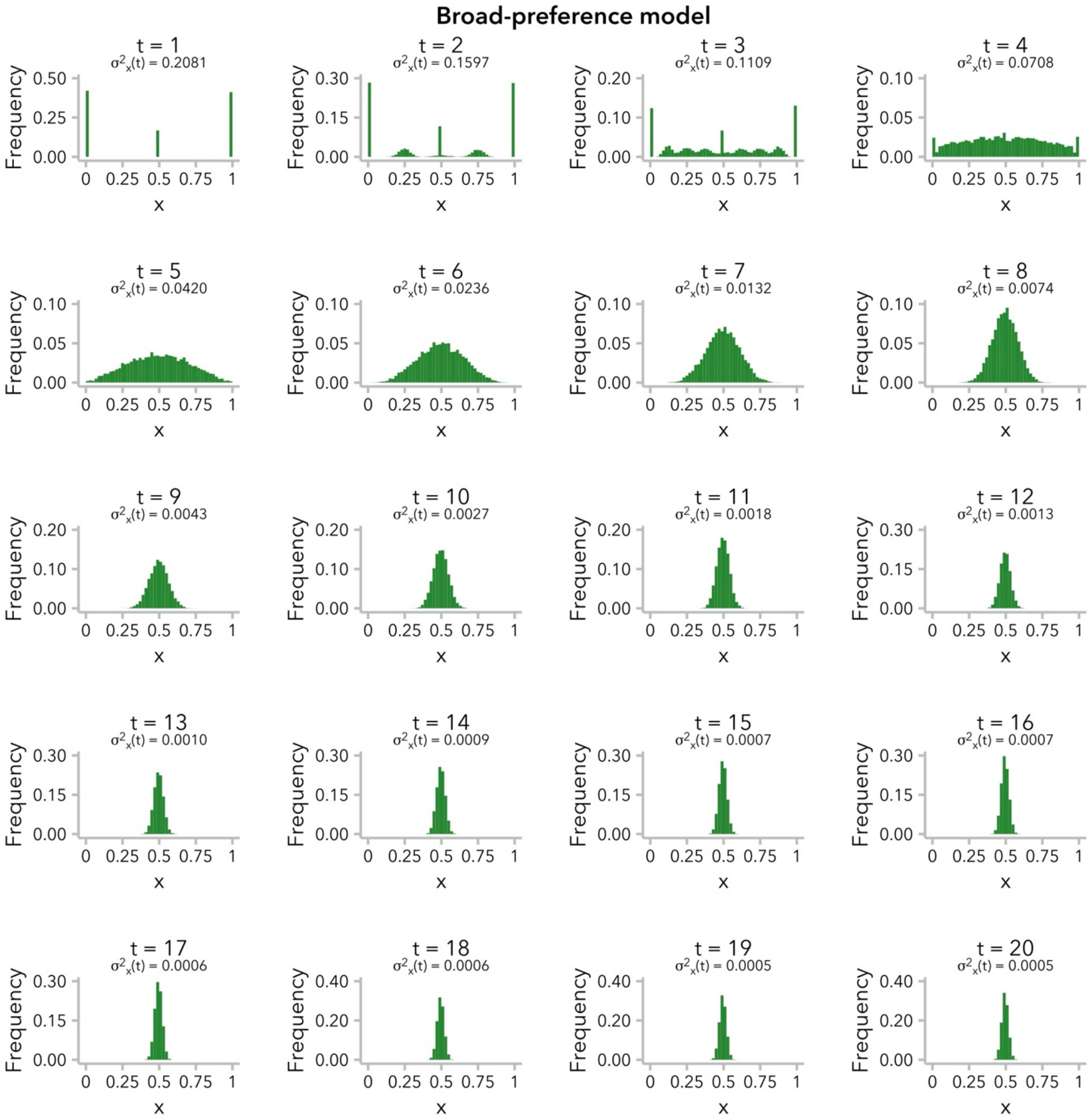


**Figure S4.** Under the broad-preference model, variance in global ancestry proportion across individuals, $\sigma_x^2(t)$, decreases as admixture proceeds. The distribution of global ancestry proportion, $x$, is plotted for the first 20 generations post-admixture for $\alpha = 5$.

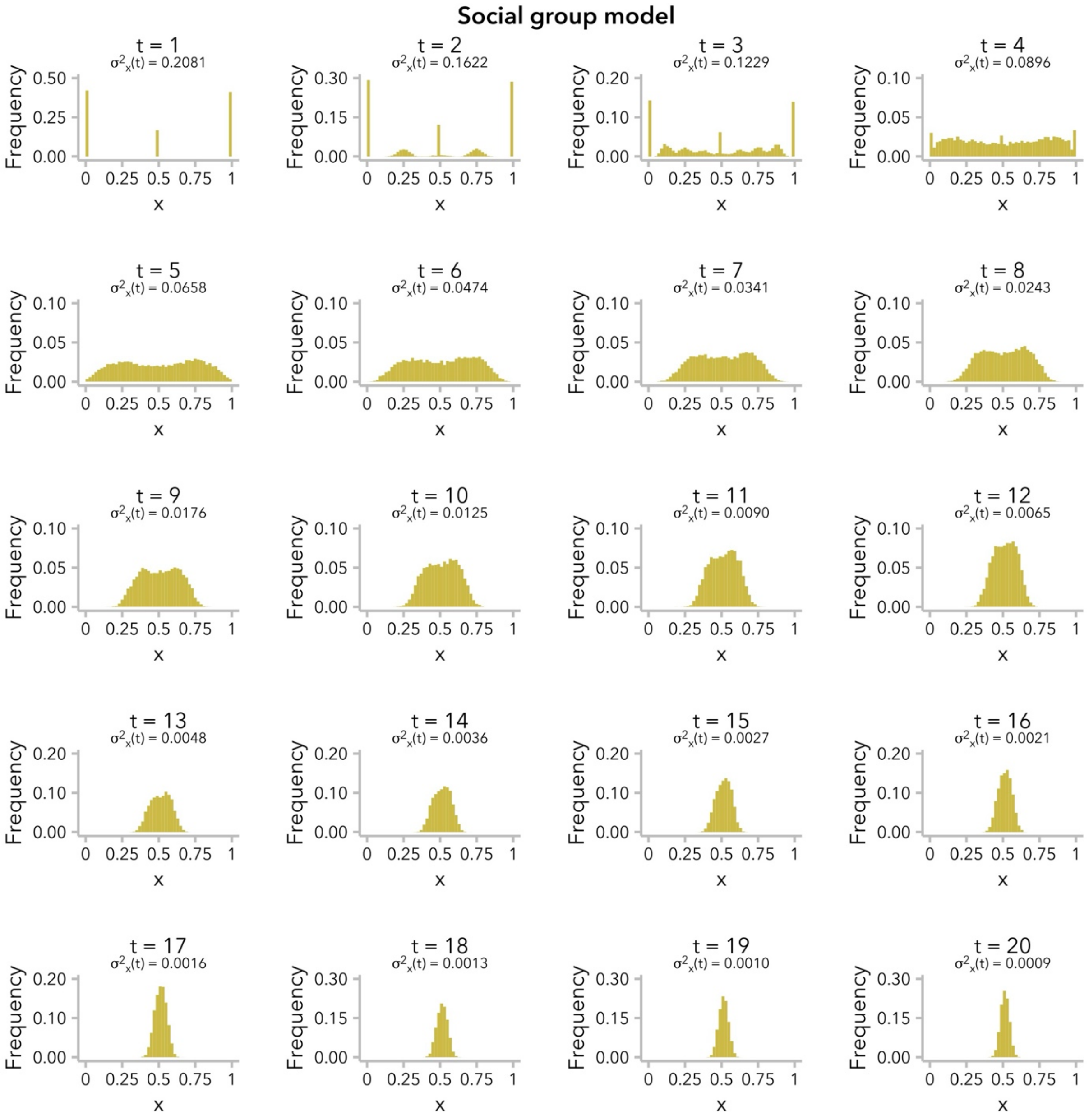


**Figure S5.** Under the social group model, variance in global ancestry proportion across individuals, $\sigma_x^2(t)$, decreases as admixture proceeds. The distribution of global ancestry proportion, $x$, is plotted for the first 20 generations post-admixture for $\alpha = 5$.

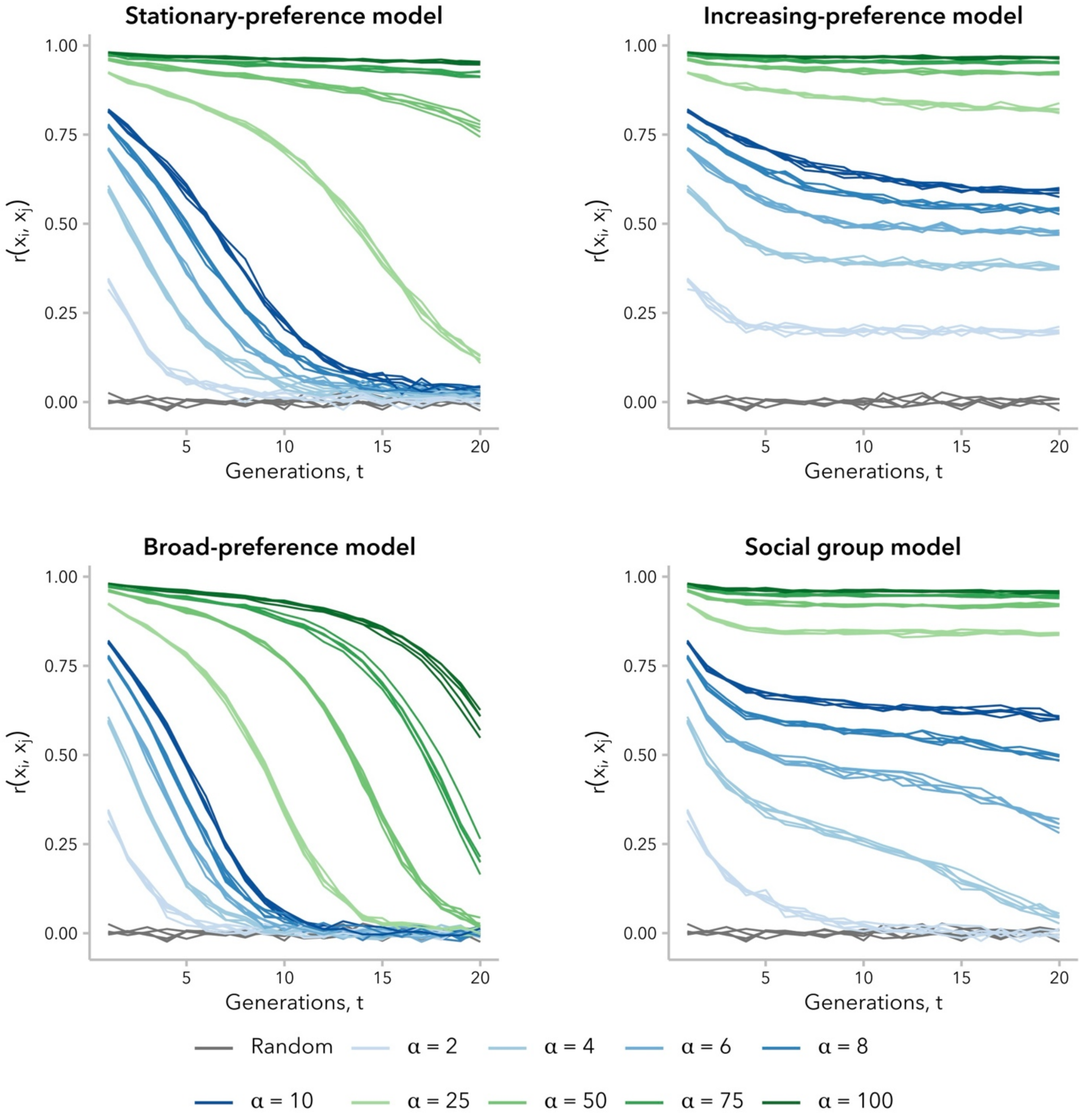


**Figure S6.** Correlation in global ancestry proportion between mates, $r(x_i, x_j)$ was not constant over time and decayed to near-zero in some simulations. For each model, higher values of $\alpha$ produced greater $r(x_i, x_j)$ in the early generations after admixture. In simulations under the stationary-preference and broad-preference models, $r(x_i, x_j)$ was near-zero at $t = 20$ generations post-admixture for $\alpha \leq 10$. The same random-mating simulations ($\alpha = 1$; gray) are reproduced in each subplot for reference. Compare to **Figure 2**.

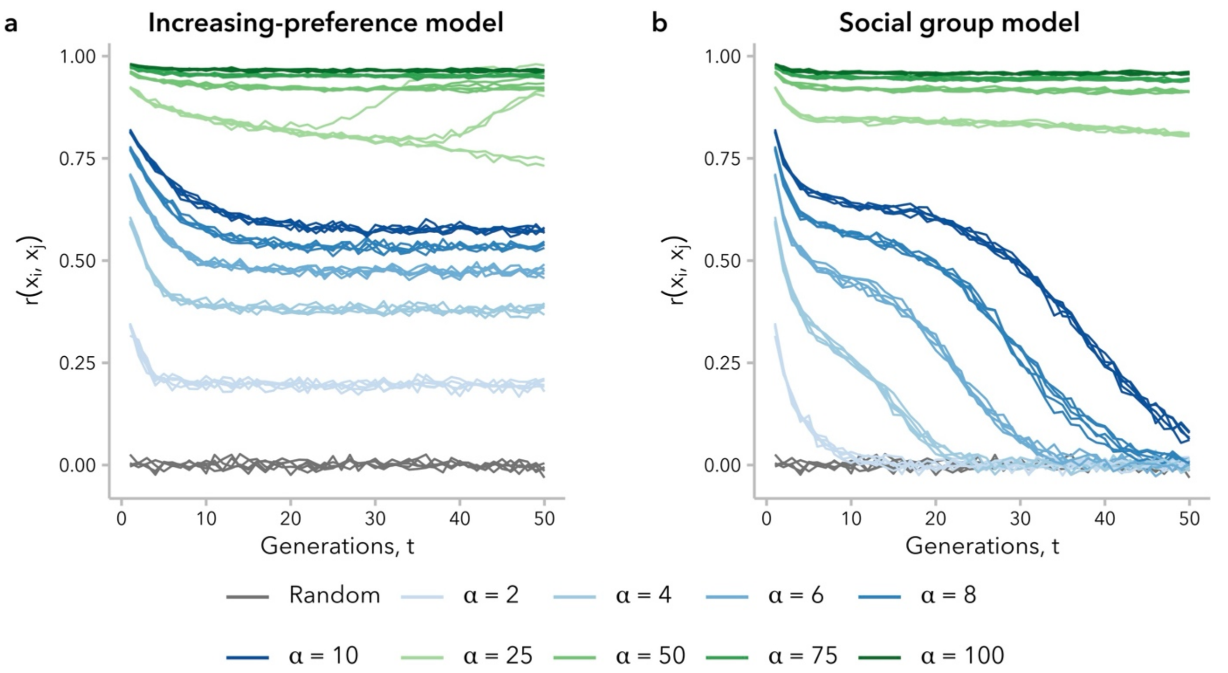


**Figure S7.** On longer time scales, the decay in the correlation in global ancestry proportion between mates, $r(x_i, x_j)$ differed between the increasing-preference and social groups models. **(a)** The value of $r(x_i, x_j)$ reached a stable non-zero plateau maintained over $t = 50$ generations post-admixture under the increasing-preference model, for all $\alpha$. **(b)** While $r(x_i, x_j) \gg 0$ for the first $t = 20$ generations post-admixture under the social group model (**Figure 2**), it approached zero on longer time scales.

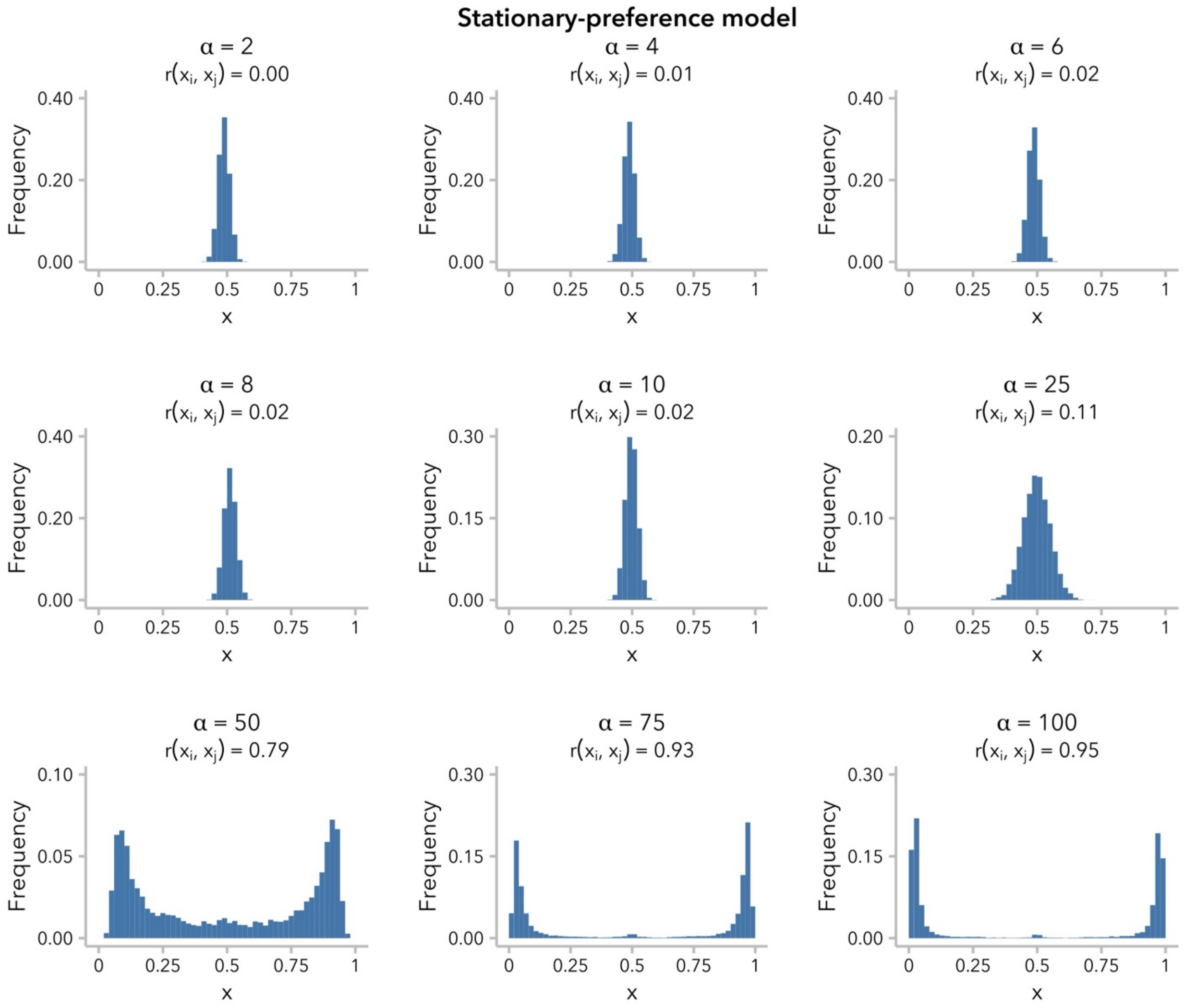


**Figure S8.** Under the stationary-preference model, a positive correlation in global ancestry proportion between mates, $r(x_i, x_j)$, was only observed at generation $t = 20$ when mate-choice bias was strong enough to largely prevent admixture ($\alpha > 10$). Each subplot displays the distribution of global ancestry proportion across individuals at $t = 20$ generations for the indicated value of $\alpha$.

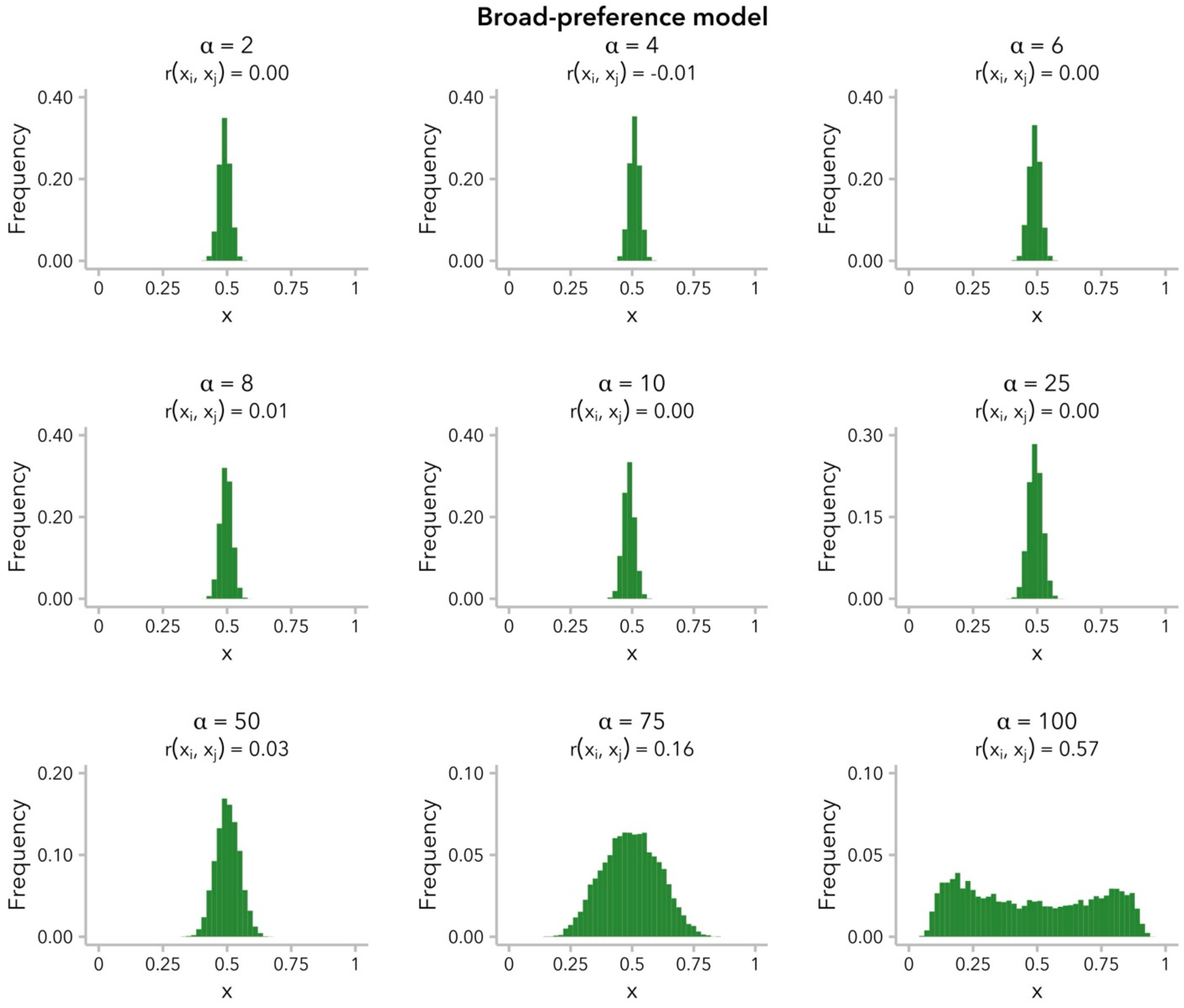


**Figure S9**. Under the broad-preference model, a positive correlation in global ancestry proportion between mates, $r(x_i, x_j)$, was only observed at generation $t = 20$ when mate-choice bias was strong enough to largely prevent admixture ($\alpha > 10$). Each subplot displays the distribution of global ancestry proportion across individuals at $t = 20$ generations for the indicated value of $\alpha$.

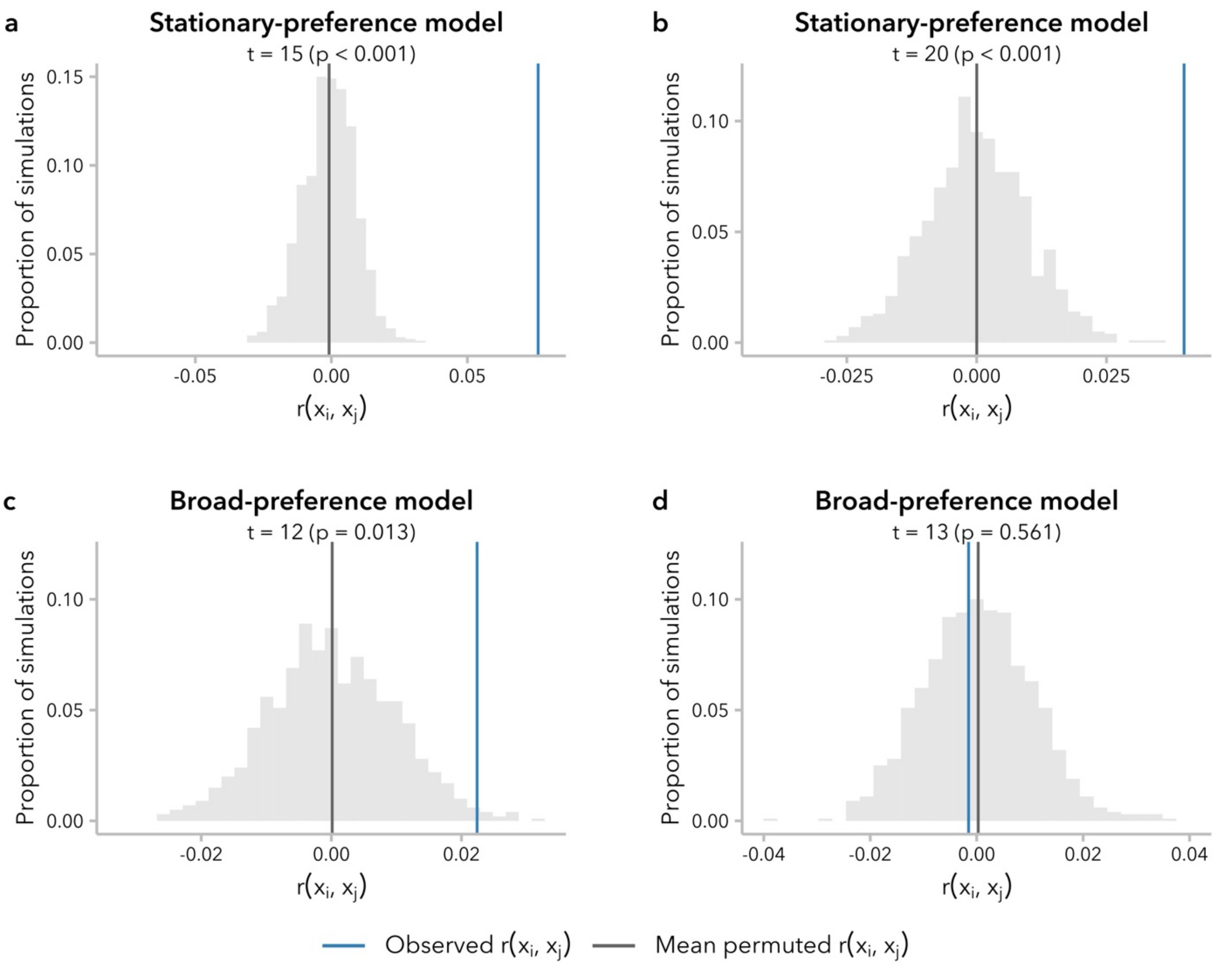


**Figure S10.** Mating can differ significantly from random even when the correlation in global ancestry proportion between mates, $r(x_i, x_j)$, is very small. **(a, b)** In simulations under the stationary-preference model, $r(x_i, x_j)$ was significantly different from 1,000 permutations across all time points for $\alpha = 10$. **(c, d)** In simulations under the broad-preference model, $r(x_i, x_j)$ was only significantly different from permutation until generation $t = 20$ for $\alpha = 10$.

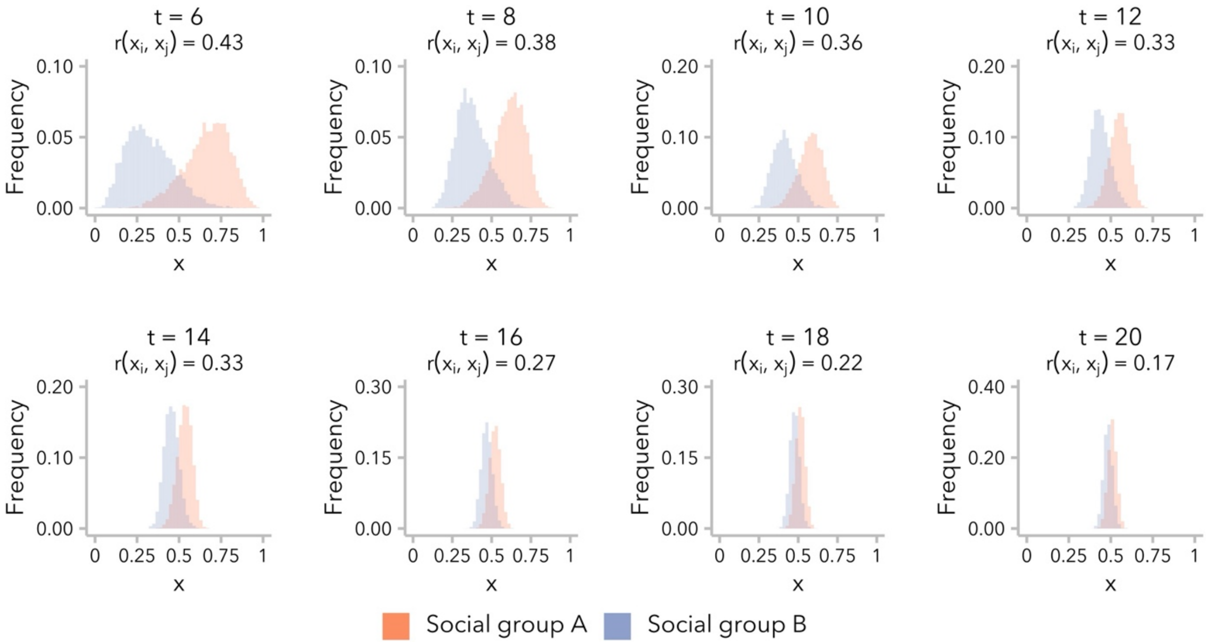


**Figure S11.** Differences in global ancestry proportion, $x$, between social groups became smaller over time in simulations run under the social group model. Despite increasing overlap in the distributions, correlation in global ancestry proportion between mates, $r\left(x_i, x_j\right) \gg 0$ at $t = 20$ generations post-admixture. A representative example simulation is shown for $\alpha = 5$.

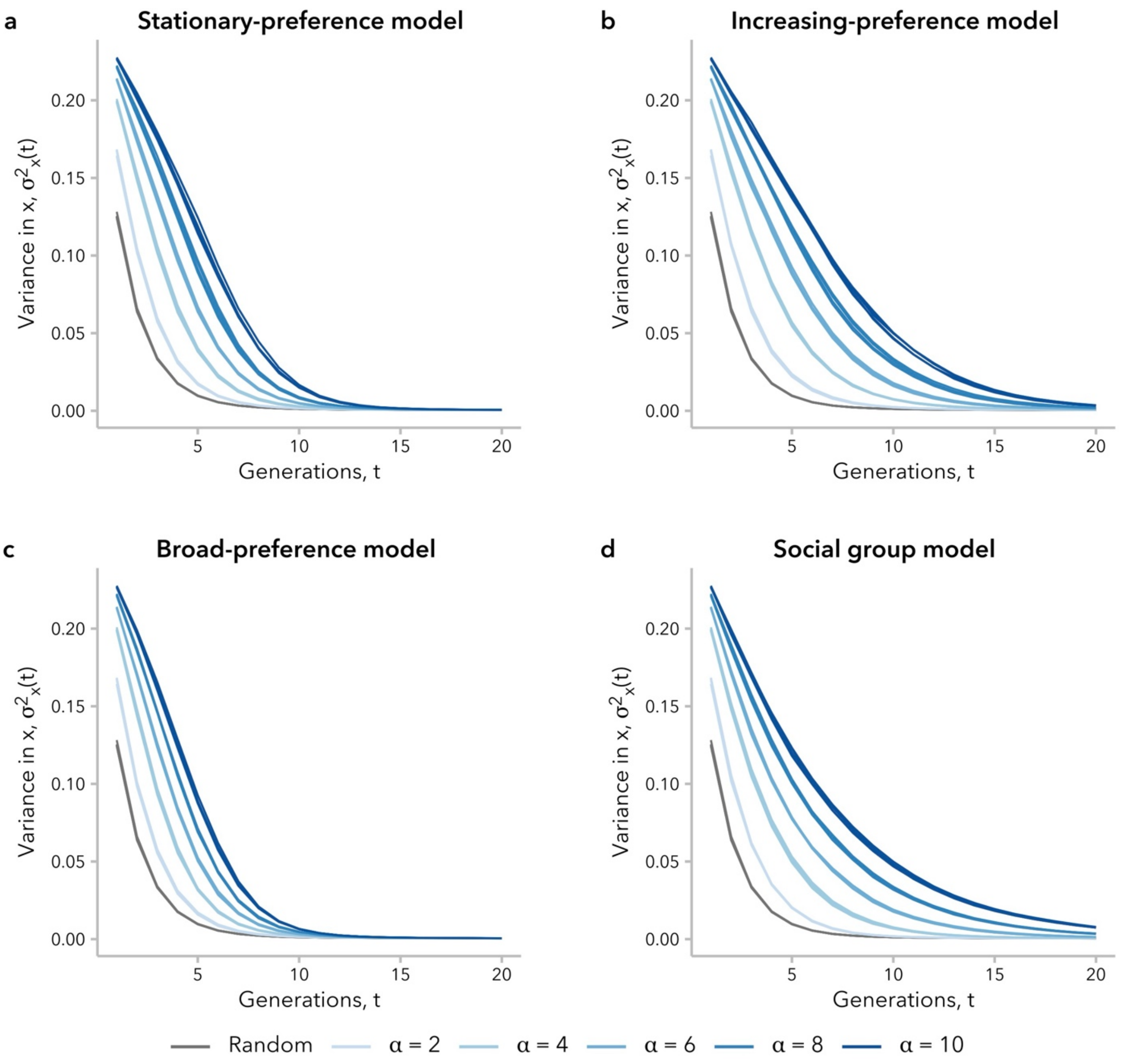


**Figure S12.** Variance in global ancestry proportion across individuals, $\sigma_x^2(t)$ decayed to zero over time for a single-pulse admixture scenario, but the decay occurred more slowly for larger values of $\alpha$. At $t = 20$ generations post-admixture, only simulations under the increasing-preference and social group models **(b, d)** had a greater $\sigma_x^2(t)$ than the random-mating control simulations.

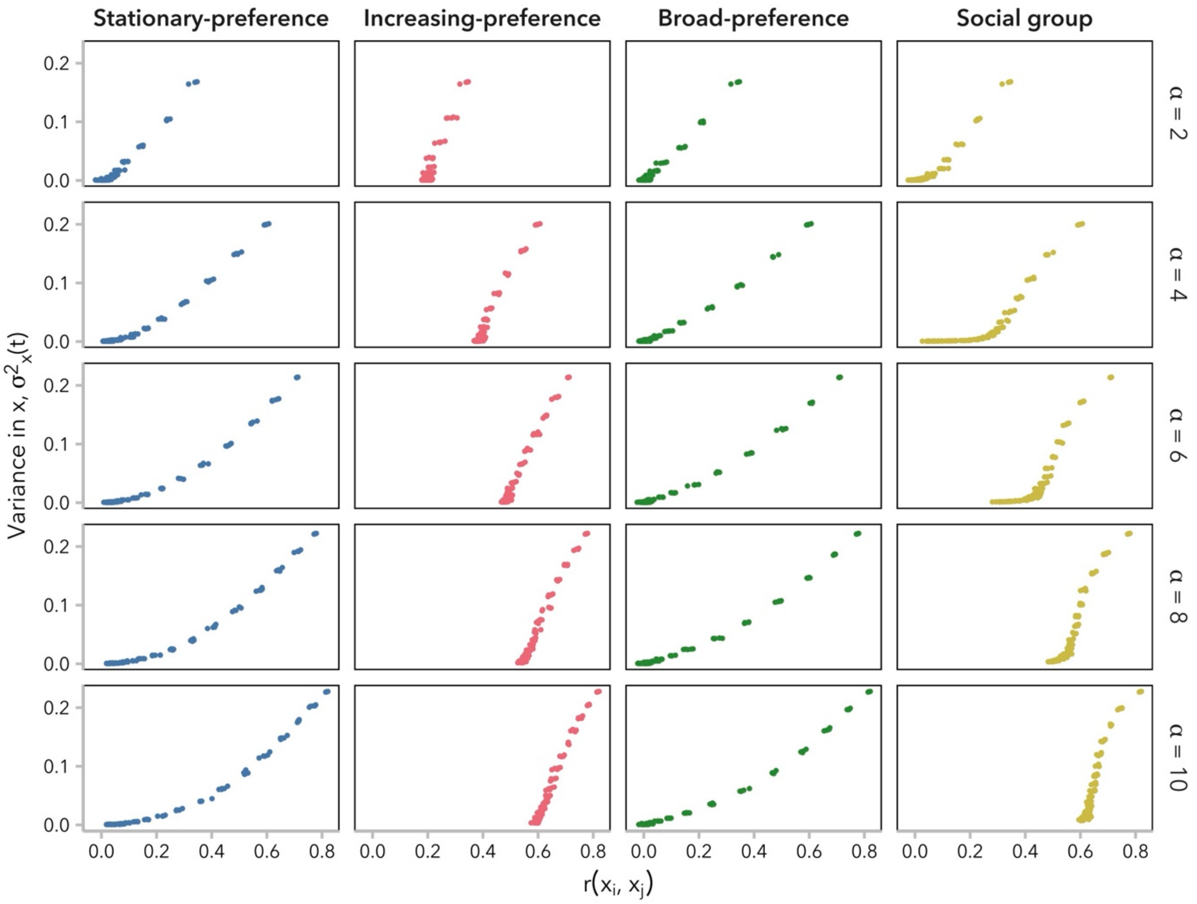


**Figure S13.** Correlation in global ancestry proportion between mates, $r(x_i, x_j)$ is positively correlated with variance in global ancestry proportion across individuals, $\sigma_x^2(t)$ for all models and values of $\alpha$. Relative to the other two models, the increasing-preference and social group models maintain $r\left(x_i, x_j\right) \gg 0$ even as $\sigma_x^2(t)$ decreases. Each dot represents one generation, $t \in [1,50]$.

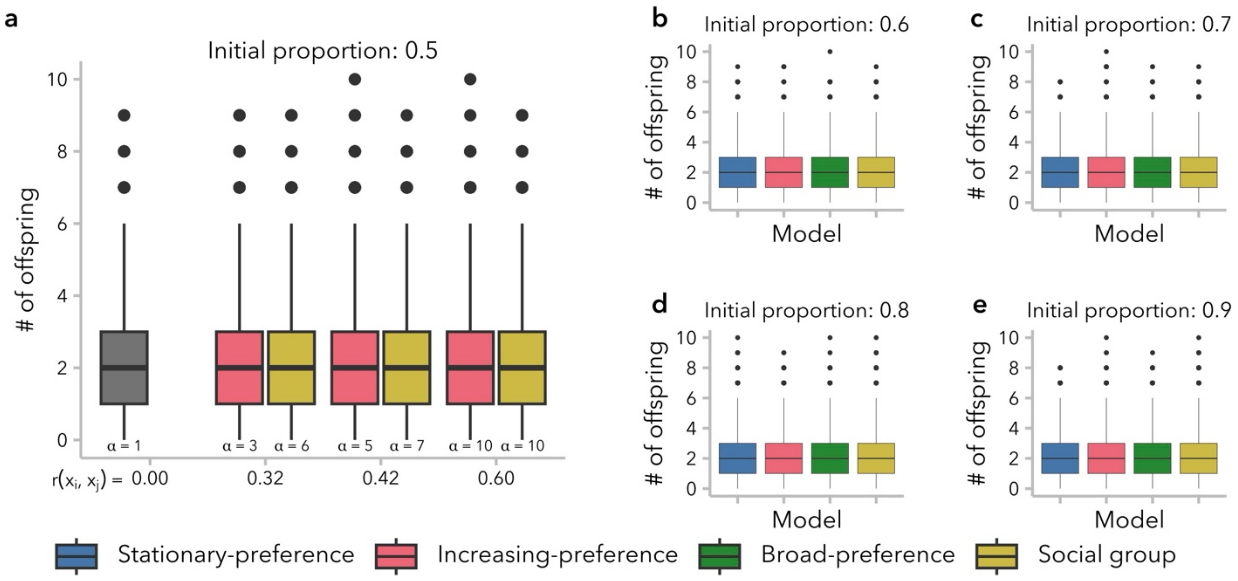


**Figure S14.** Biased mating did not impact mean reproductive success at $t = 20$ generations across models, values of $\alpha$, or the initial admixture contributions. **(a)** Three pairs of simulations, matched for the correlation in global ancestry proportion between mates, $r(x_i, x_j)$, are shown at generation $t = 20$. Over 85% individuals contributed 3 or fewer offspring to the next generation. **(b-e)** Mean reproductive success did not change with increasing contribution from source population 1 to the founding of the admixed population. One replicate simulation is shown for each model at $t = 20$ generations for $\alpha = 10$.

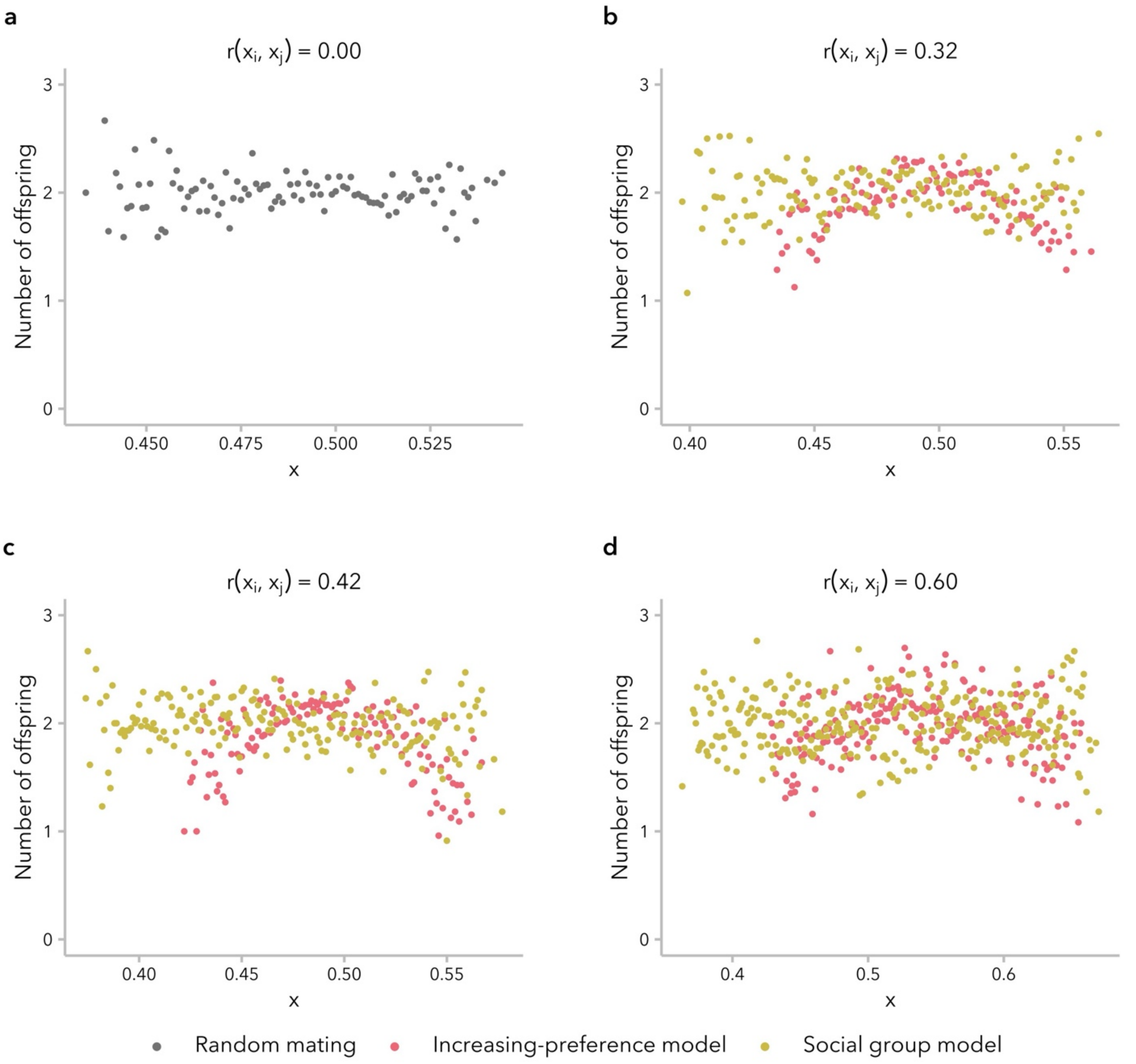


**Figure S15.** The effects of global ancestry proportion, $x$, on reproductive success was negligible, regardless of the correlation in global ancestry proportion between mates, $r(x_i, x_j)$. Three pairs of simulations, matched for the $r(x_i, x_j)$ are shown at generation $t = 20$. Each dot represents the mean number of offspring in a bin 0.001 $x$ units. Bins containing with fewer than 10 individuals are not shown.

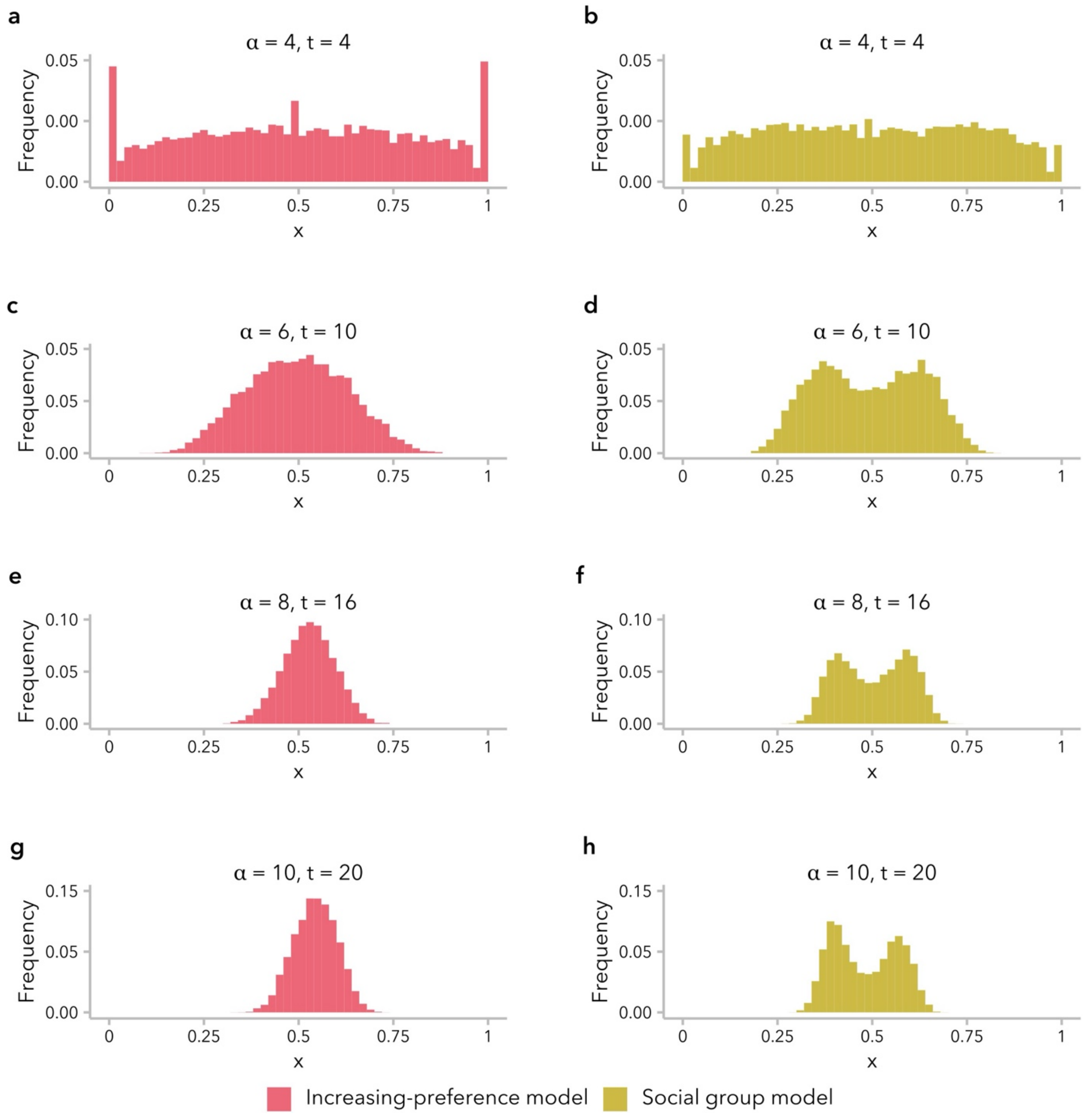


**Figure S16.** The admixture process results in a transition in the shape of the distribution of global ancestry proportion, $x$, from bimodal to unimodal. Holding $\alpha$ constant, this transition occurred later in simulations run under the social group model (right column: **b**, **d**, **f**, **h**), relative to the increasing-preference model (left column: **a**, **c**, **e**, **g**). Four representative examples are shown: each row compares the two models for the same $\alpha$ and generation $t$.

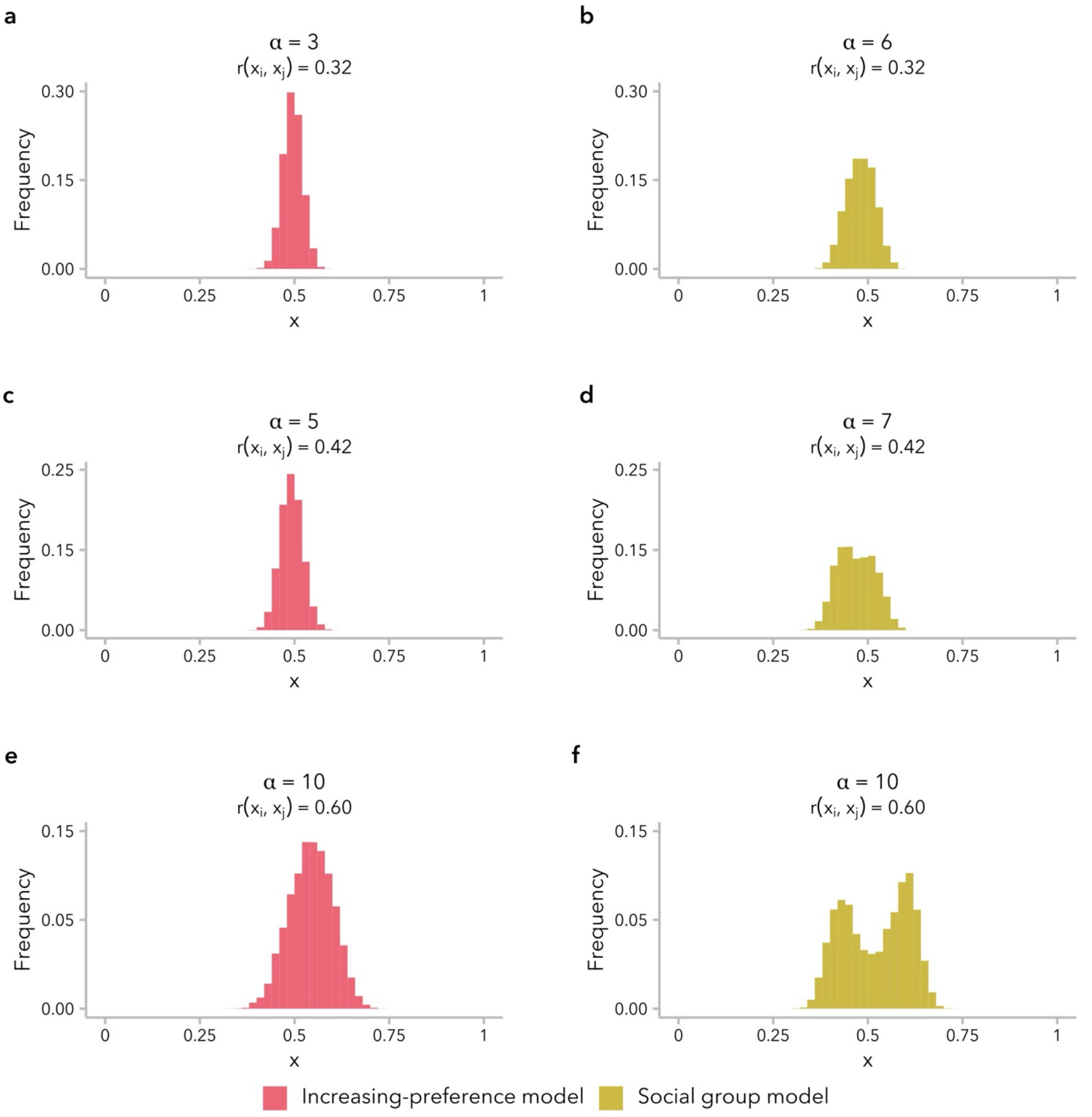


**Figure S17.** The distribution of global ancestry proportion, $x$, was unimodal at $t = 20$ generations post-admixture in simulations under the increasing-preference model (left column: **a**, **c**, **e**) and remained bimodal in simulations under the social group model (right column: **b**, **d**, **f**). The bimodal shape of the distribution was more pronounced for larger values of $\alpha$. Three examples are shown, comparing simulations with similar correlation in global ancestry between mates, $r(x_i, x_j)$ at generation $t = 20$.

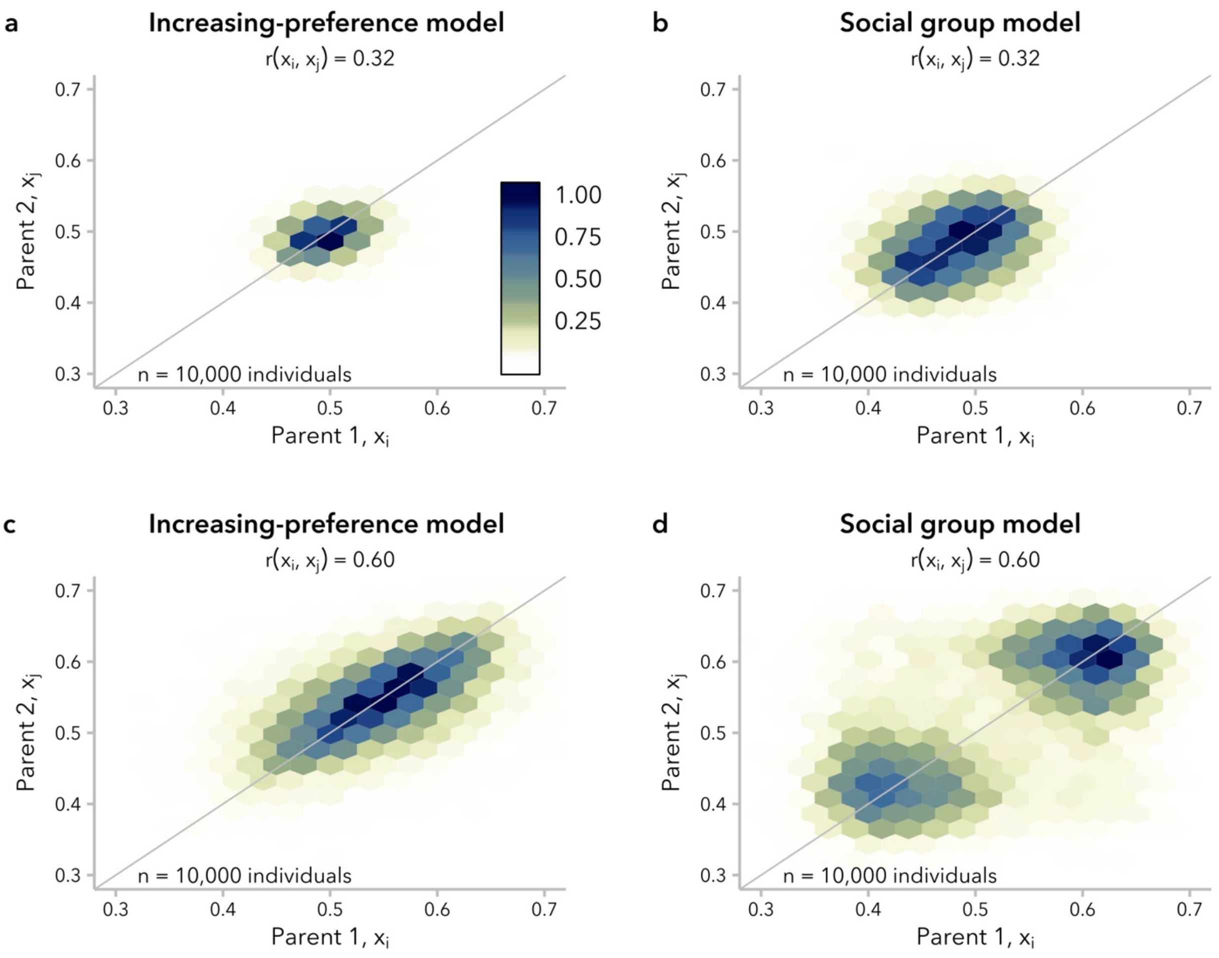


**Figure S18.** Differences in mating structure between simulations under the increasing-preference **(a, c)** and social group **(b, d)** models were more pronounced when the correlation in global ancestry proportion between mates, $r(x_i, x_j)$ was larger. The parent pairs of all individuals in generation $t = 20$ are shown. Each hexagon represents a bin of 0.025 global ancestry proportion units, with color encoding the scaled density ($\max = 1$) of mating pairs in each bin. **(a)** $\alpha = 3$ **(b)** $\alpha = 6$ **(c)** $\alpha = 10$ **(d)** $\alpha = 10$. Compare to **Figure 4a.**

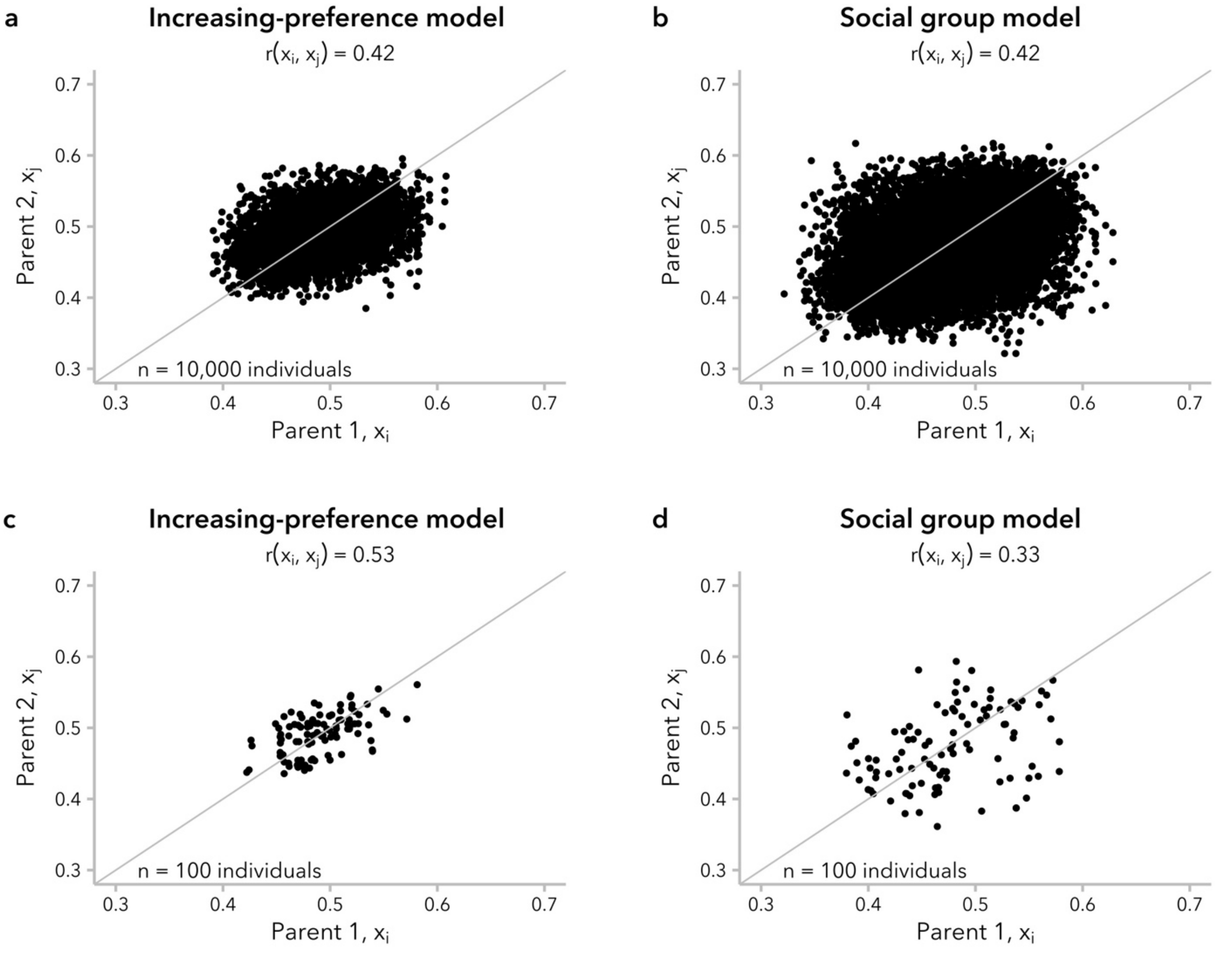


**Figure S19.** Simulations under the increasing-preference ($\alpha = 5$) and social group ($\alpha = 7$) models produced the same correlation in global ancestry proportion, $r(x_i, x_j)$, with different underlying mating structure. Differences in the underlying structure were difficult to ascertain when visualized as a dot-plot. Each dot represents the parent pair of an individual in generation $t = 20$. **(a, b)** All mating pairs ($n = 10{,}000$). **(c, d)** A random sample of 100 mating pairs ($n = 100$).

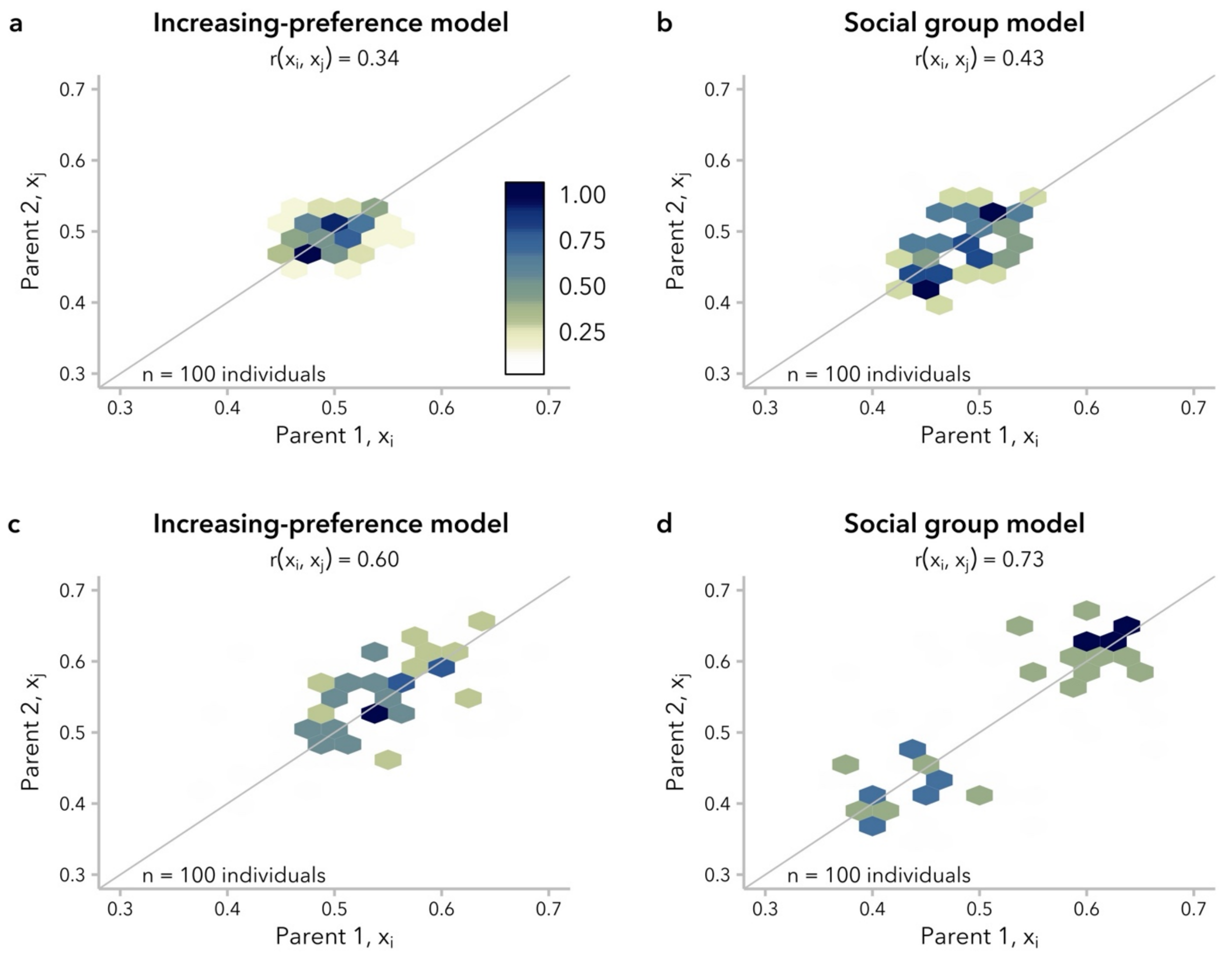


**Figure S20.** Even with a relatively small sample size ($n = 100$), hexagonal bin plots can suggest differences between the increasing-preference (**a, c)** and social group **(b, d)** models. Differences in the structure of mating pairs is more apparent for simulations with greater correlation in global ancestry proportion between mates, $r(x_i, x_j)$. **(a)** $\alpha = 3$ **(b)** $\alpha = 6$ **(c)** $\alpha = 10$ **(d)** $\alpha = 10$. Compare to **Figure 3c**.

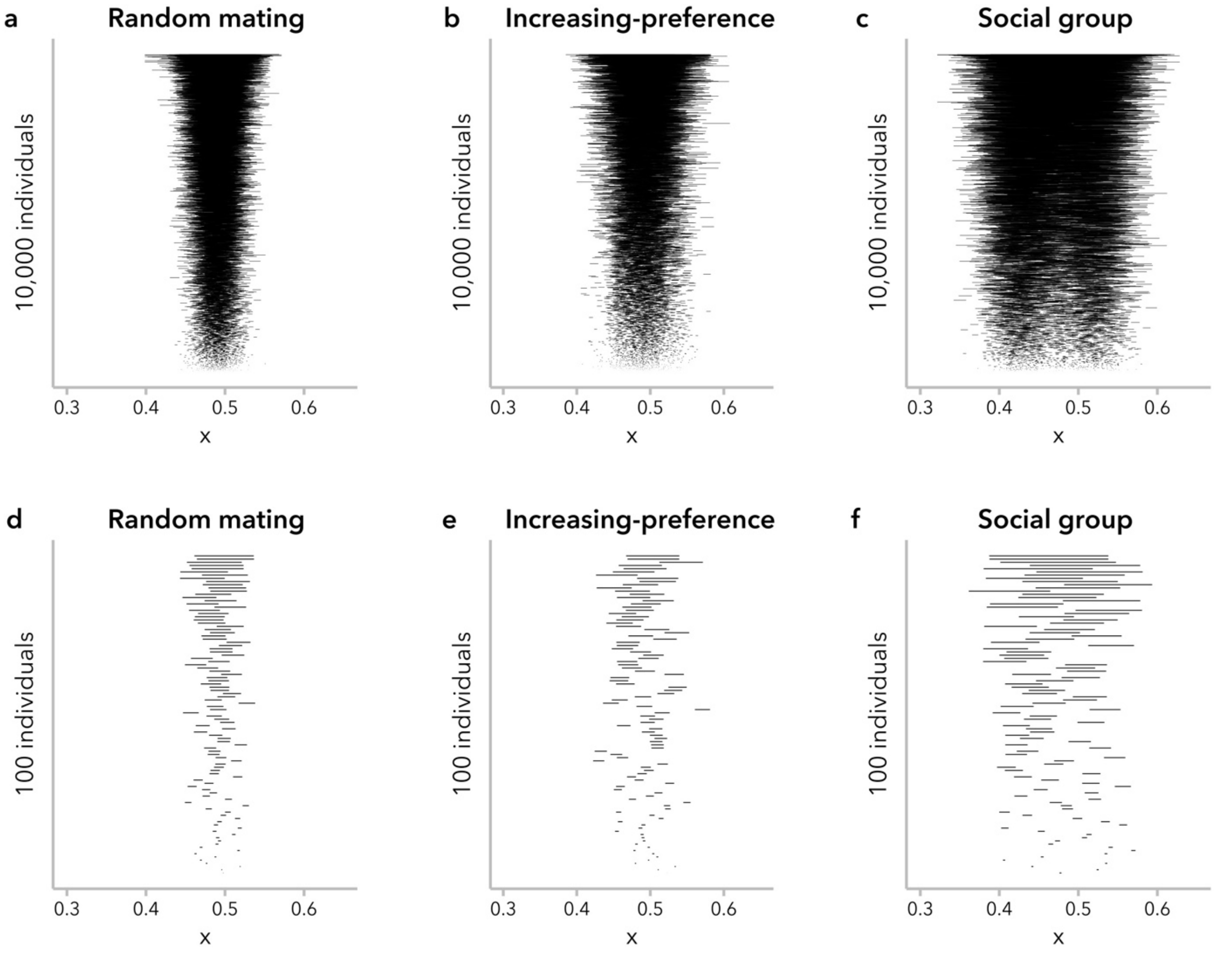


**Figure S21.** In addition to hexagonal bin plots, hurricane plots visualizing the absolute difference in global ancestry proportion between mate pairs, $\Delta_x = |x_i - x_j|$, can suggest differences in mating structure between the increasing-preference and social group models. Representative examples are shown for **(a)** random mating ($\alpha = 1$), **(b)** the increasing-preference model ($\alpha = 10$), and **(c)** the social group model ($\alpha = 10$). Each line represents one individual in generation $t = 20$, connecting the global ancestry proportion, $x$, of the two parents. **(d-f)** a subset of 100 individuals are sampled from the population.

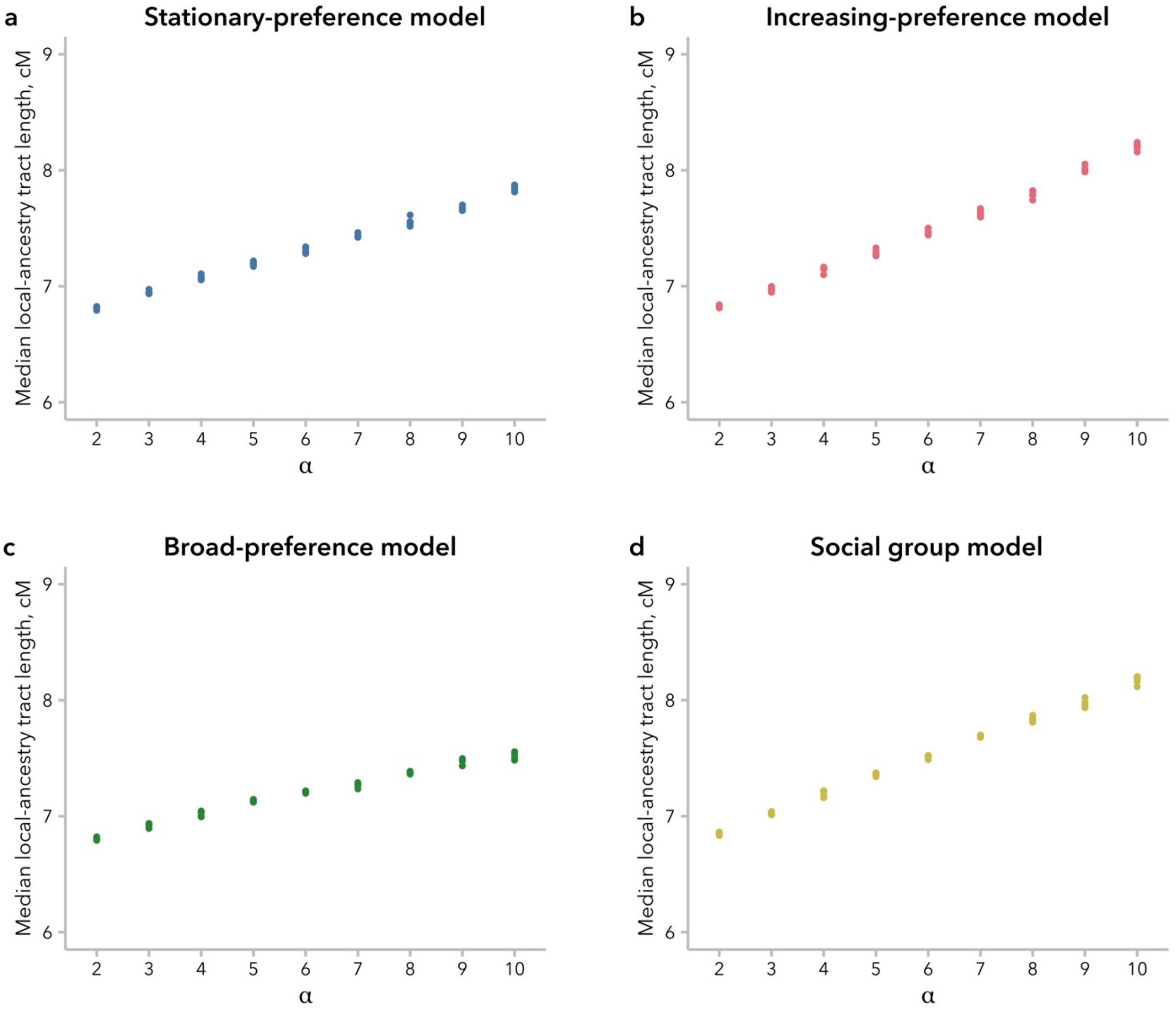


**Figure S22.** Comparing within each model, stronger mate-choice bias (larger values of $\alpha$) consistently correspond to longer median length of local-ancestry tracts. Five replicate simulations are shown for each value of $\alpha$.

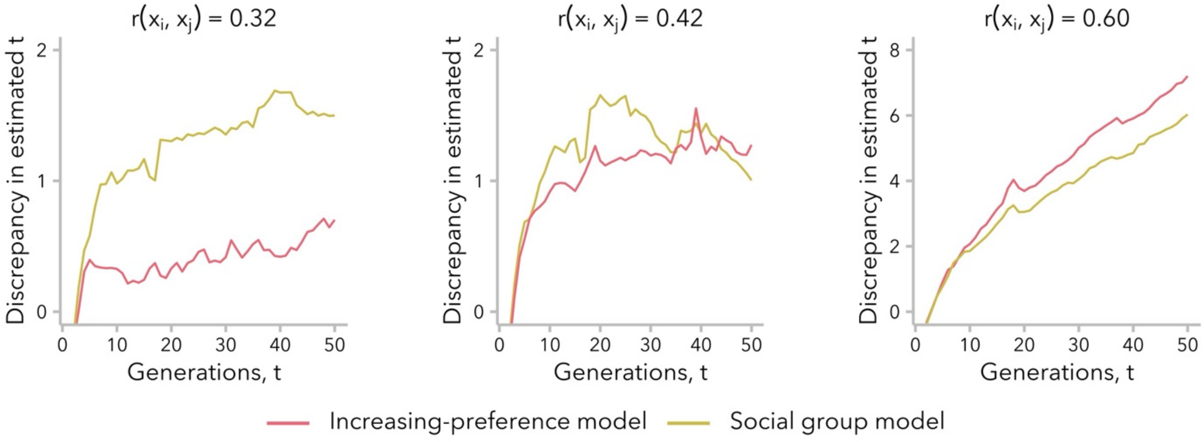


**Figure S23.** Underestimation of the time since admixture is more strongly influenced by earlier generations. In simulations under either the increasing-preference or social group model, the discrepancy between the estimated time since admixture and the true time since admixture did not increase linearly over time, and in some cases appeared to reach a plateau. Each line represents a single simulation replicate, matched for the correlation in global ancestry proportion between mates, $r(x_i, x_j)$.

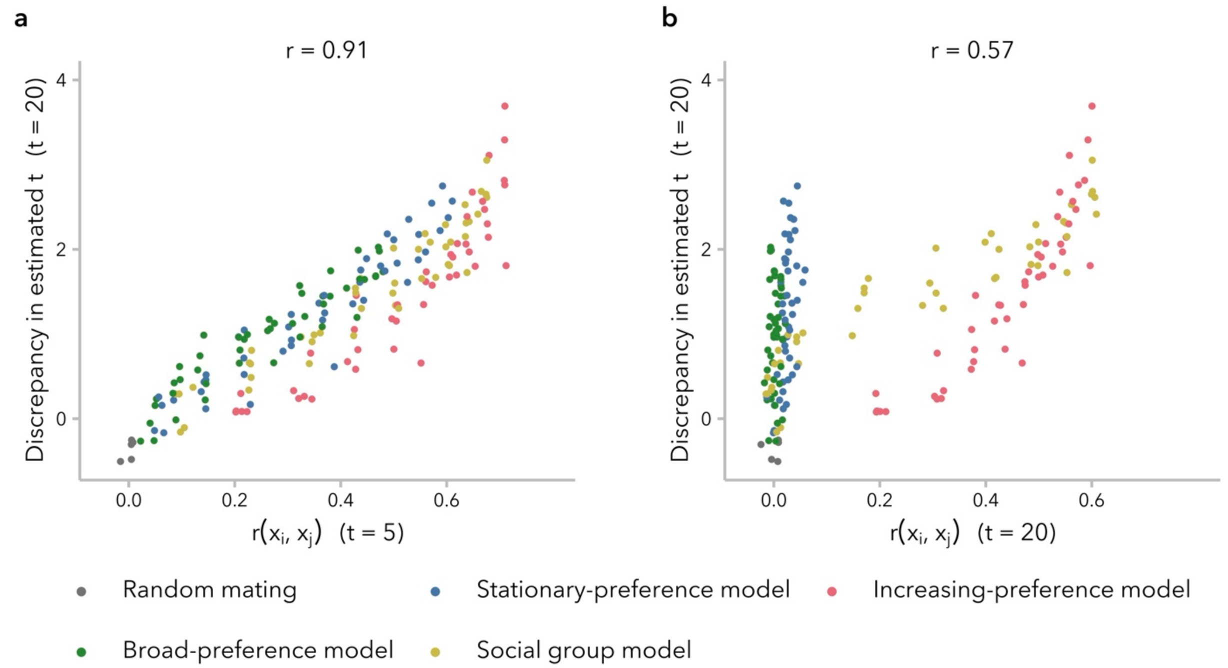


**Figure S24.** Underestimation of the time since admixture is more strongly influenced by earlier generations. The correlation in global ancestry proportion between mates, $r(x_i, x_j)$, at generation $t = 5$ more strongly influenced the discrepancy in estimated time since admixture calculated for generation $t = 20$ (**a**) than did the value of $r(x_i, x_j)$ at $t = 20$ (**b**).

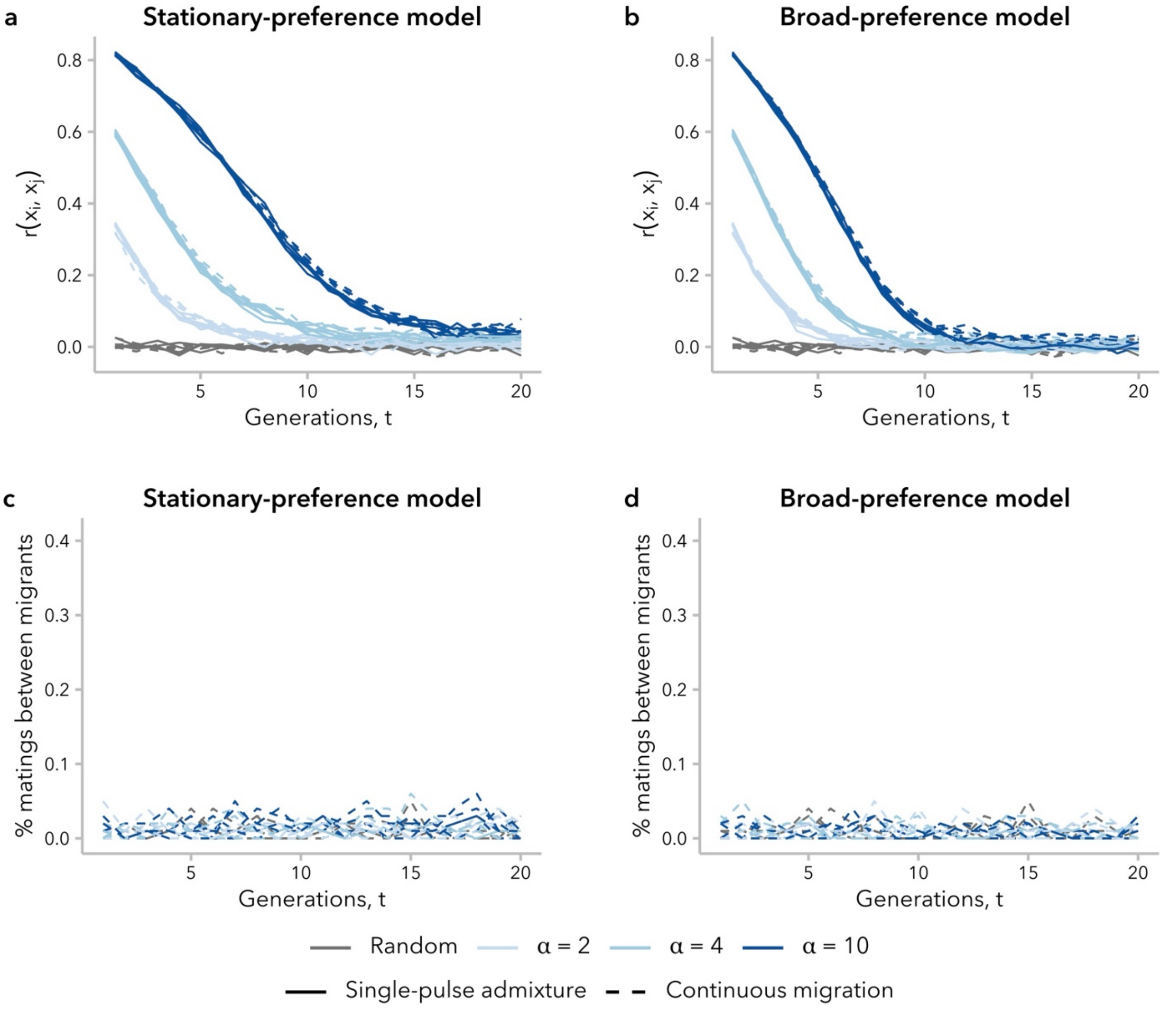


**Figure S25.** Continuous migration did not affect the correlation in global ancestry proportion between mates, $r(x_i, x_j)$, under the stationary-preference and broad-preference models. **(a, b)** The value of $r(x_i, x_j)$ did not differ meaningfully between simulations with and without migration, matched for $\alpha$. **(c, d)** Under both models, mating events involving two migrants did not occur more often than expected by chance.

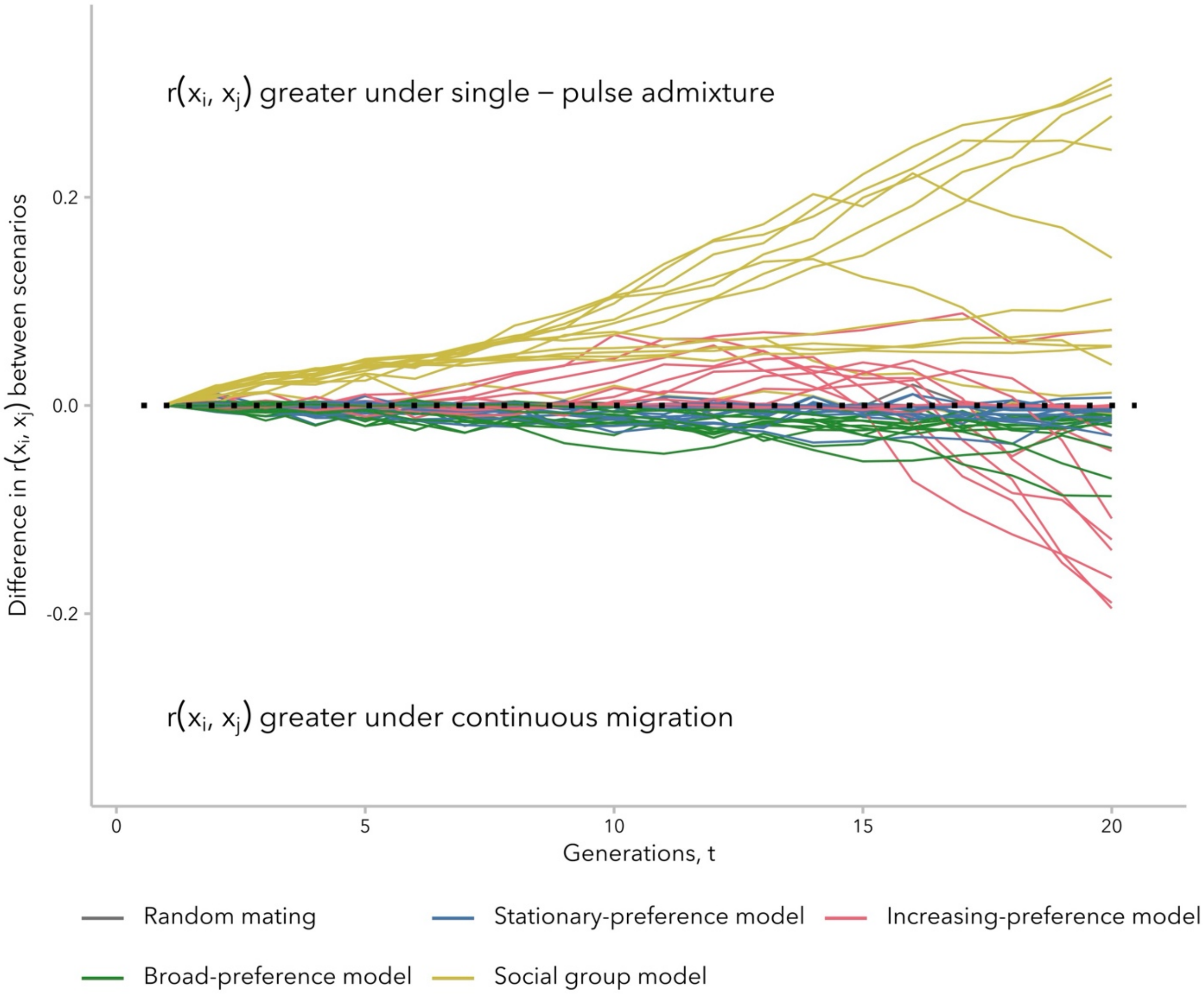


**Figure S26.** The effects of continuous migration differed between mate-choice models. The introduction of continuous migration tended to increase the correlation in global ancestry proportion between mates, $r(x_i, x_j)$, under the increasing-preference model, relative to a scenario of pulse-admixture. The opposite trend was observed under the social group model. The value of $r(x_i, x_j)$ was generally unaffected by migration under the stationary-preference and broad-preference models.

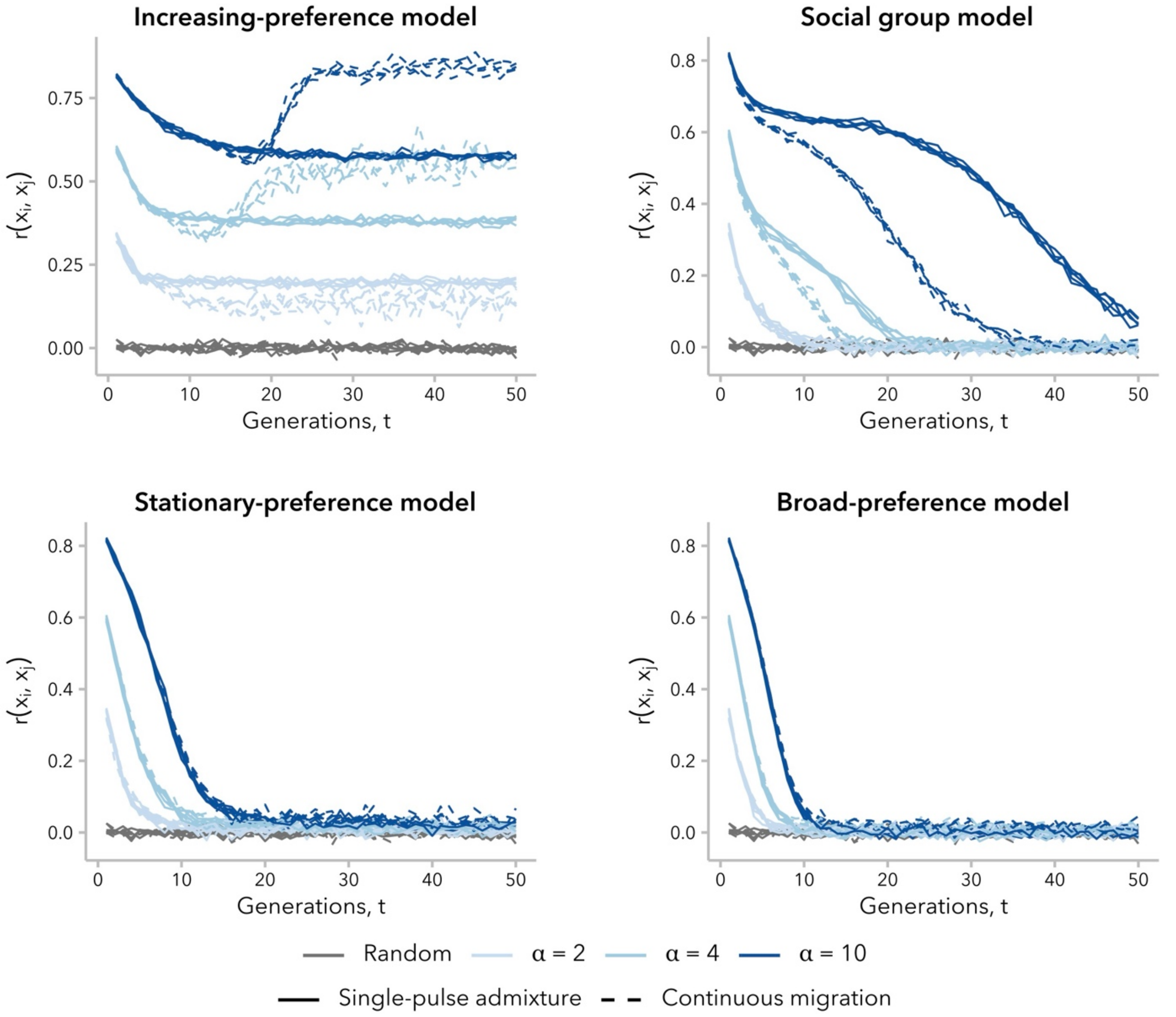


**Figure S27.** The disparate impacts of continuous migration between models were consistent on longer time scales. The correlation in global ancestry proportion between mates, $r(x_i, x_j)$, with and without continuous migration is shown under each model for 50 generations post-admixture.
